# Millisecond-scale motor coding precedes sensorimotor learning in songbirds

**DOI:** 10.1101/2024.09.27.615500

**Authors:** Leila May M. Pascual, Aanya Vusirikala, Ilya M. Nemenman, Samuel J. Sober, Michael Pasek

## Abstract

The changes in neural activity that underlie motor skill acquisition during development are unknown. Juvenile songbirds learn their songs by an imitative process, and after learning adults use a millisecond-precise spike timing code to control vocal acoustics. Current theories suggest that developmental changes in neural firing rates, rather than precisely timed spike patterns, underlie the emergence of learning. Here we tested the hypothesis that songbirds transition from a rate-based to a spike-timing-based motor code during the process of vocal development. To do so, we recorded vocal motor neurons across development as individual Bengalese finches learned their songs. Contrary to our hypothesis, we found that despite dramatic changes in firing statistics during development, millisecond-scale spike pattern codes for vocal acoustics are present throughout all stages of vocal development. Furthermore, firing rate fluctuations are no more predictive of song output in young learners than in expert adults. The dramatic changes in spiking statistics observed during song learning therefore do not reflect a developmental change in the timescale of motor coding, but instead signals the selection of a particular subset of precisely timed spike patterns. We speculate that such patterns are favored because they most reliably modulate behavior.

## Introduction

Motor skills are acquired through practice until they are performed at high levels of precision. During this process of motor learning, the brain uses sensory feedback to refine the neural commands transmitted to muscles to shape behavior [1–4]. To investigate the changes in brain activity that support this learning process, prior studies have quantified fluctuations in individual neurons’ “spike rates” (the total number of spikes fired across hundreds of milliseconds). Such studies have consistently observed that spike rate variability decreases as behavior becomes more stereotyped during motor skill learning [5–10]. These findings might mean that the mapping between neural activity and behavior (the “motor code”) relies on variations in spike rate early in life, and that the reduction in behavioral variability results from the observed reduction in spike rate variability. However, whether the motor code is based on spike rates (rather than precise spike timing at finer timescales) has only been tested in adult animals in a small number of species [11–14], and to our knowledge developmental changes in the timescale of the motor code have never been directly tested.

The developing songbird provides a particularly powerful system in which to examine motor skill development. During song learning, juvenile songbirds develop new vocal gestures called “syllables” (Fig. 1) which are acoustically refined until they become “crystallized” by adulthood [15–17]. Previous findings in juvenile songbirds have paralleled results in other species: the variability of cortical firing rates in juvenile zebra finches decreases over the same period that variability in vocal behavior decreases across learning [10]. Furthermore, our prior work has shown that adult songbirds use millisecond-scale variations in millisecond-precise spike timing patterns to control behavior, including patterns that exploit nonlinearities in muscular control [11, 18, 19]. Together, these findings suggest that changes in neural activity during song learning might reflect the nervous system shifting from rate-based control of song acoustics (as juveniles) to the spike timing-based control we previously demonstrated in adult songbirds [11].

**Fig. 1.**
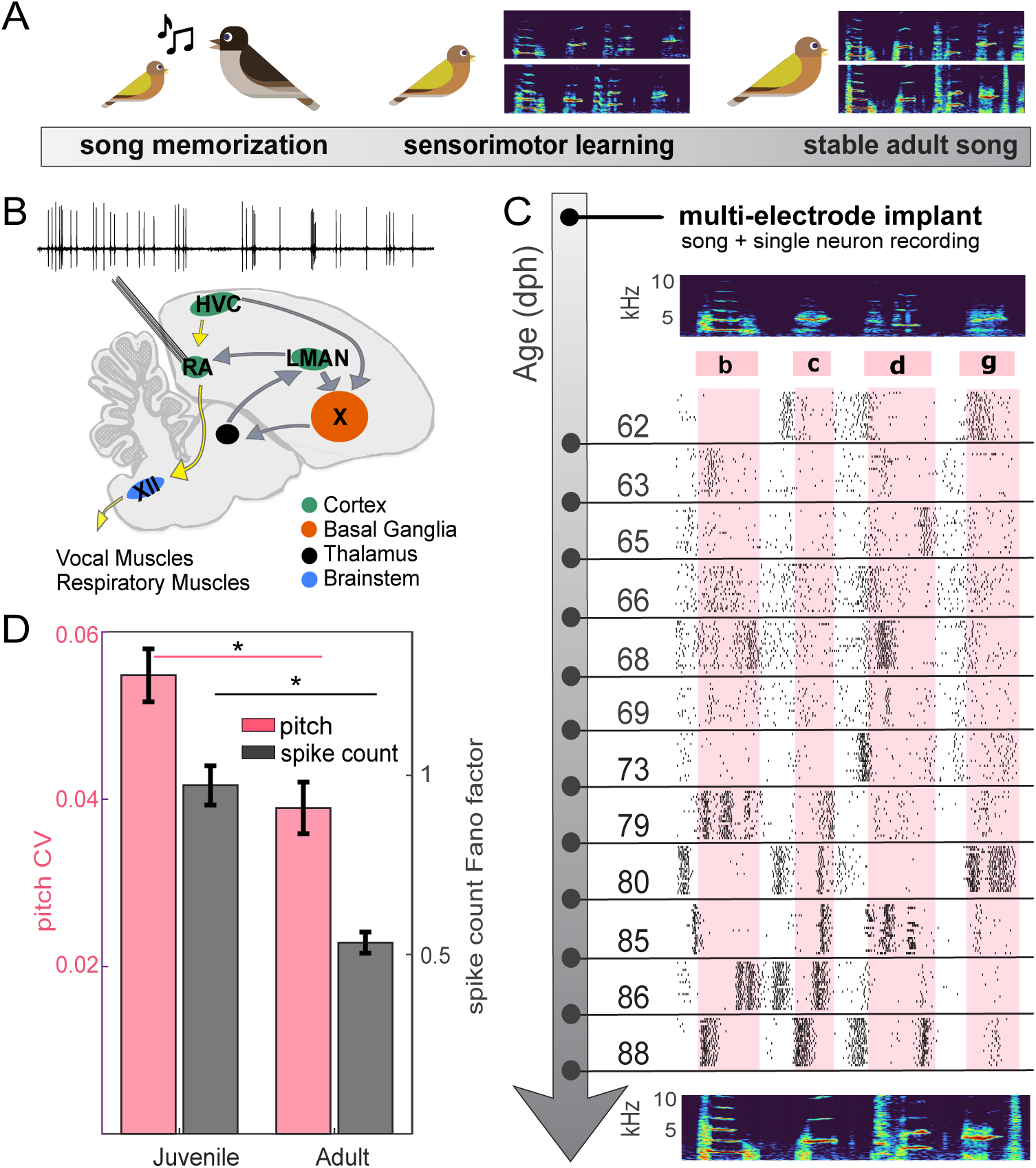
Recordings across song learning in juvenile Bengalese finches reveal changes in gross statistics of neural spiking. **A)** A juvenile songbird begins learning its song with early life exposure to a tutor’s song and continues with sensorimotor practice across development. Spectrograms illustrate high variability in juvenile song renditions (middle) compared to the stereotyped adult song (right). **B)** Recordings targeted projection neurons in vocal motor nucleus RA (representative voltage trace of an RA neuron recorded at 68 days post hatch (dph) is shown). RA receives synaptic input from HVC and LMAN and innervates brainstem vocal motor neurons (nXIIts) via the song motor pathway (yellow arrows) [24]. **C)** Longitudinal recordings from a multi-electrode implant: spectrograms show a representative motif (repeated sequence of syllables b,c,d,g) during early (top) and late (bottom) learning. Spike rasters display 20 trials from RA neurons at indicated ages (dph); shading denotes mean syllable duration. Spike trains are linearly time-warped to the mean duration of syllables and inter-syllable pauses for visualization only (see Spike train visualization and analysis; note: quantitative analyses used unwarped spike times). **D)** Pitch coefficient of variation (CV, pink, juvenile *n* = 68, adult *n* = 37) and spike count Fano factor (gray, juvenile *n* = 101, adult *n* = 47) significantly decrease from juvenile to adulthood (two-sample KS tests, *p *<* 0.05). Error bars: mean *±* SEM.

Here we ask whether the use of millisecond-precise spike patterns to control song behavior is present from early in vocal development or emerges with time and practice. Importantly, the force-producing properties of songbird vocal muscles change significantly during development [20, 21], suggesting that the mapping between neural activity and song acoustics might similarly change between a bird’s first vocalizations and its fully learned adult song. We therefore hypothesized that the previously-observed developmental reduction in spike rate variability [22] reflects, and exploits, such a biomechanical change, shifting patterns of neural variability from slower variations in spike count to millisecond-scale timing patterns as the vocal muscles mature.

We tested this hypothesis by asking whether song learning is accompanied by a change in the timescale with which motor cortical activity encodes vocal acoustics. We combined acoustic recordings of song at different developmental time points with electrophysiologcal recordings of single neurons and quantified learning-related changes in the neural code for song. We recorded from neurons in vocal motor cortex—the robust nucleus of the arcopallium (RA)—at different ages in juvenile songbirds and examined the activity patterns preceding individual syllables (Fig. 1B-C). We then used information-theoretic analyses to determine the timescale at which neural activity best predicts vocal acoustics, allowing us to test our hypothesis that unlike the millisecond-precise spike timing code used by adult songbirds [11], a spike rate code predominates earlier in development. We found that RA neurons in the Bengalese finch display significant reductions in rate variability and increases in firing sparseness across the sensorimotor learning period, consistent with previous findings in juvenile zebra finches [10]. We then quantified whether and how the temporal resolution of the motor code changes across development by estimating the mutual information (MI) between syllable acoustics and spiking patterns at different temporal resolutions [11, 23]. Contrary to our hypothesis that the timescale of vocal motor coding changes from slower variations to millisecond precision over the course of learning, our analysis revealed a consistently precise motor code with millisecond-level structure in both juvenile birds (even at the earliest ages recorded) and adult animals.

Our results suggest that vocal learning does not reflect a shift from a less precise, rate-based code to a more temporally precise one, but rather a refinement in the “vocabulary” of particular millisecond-scale spike patterns that produce desired acoustic output. Moreover, our work suggests that studies that examine only firing rate statistics as correlates of learning may overlook more stable (and consequential) properties of neural coding based on precise multi-spike timing patterns.

## Results

To examine how cortical activity evolves across developmental skill acquisition, we recorded single-unit RA activity and song acoustics throughout the sensorimotor learning period in juvenile Bengalese finches (Fig. 1A-B). As described previously [25], we found that the acoustic variability of song syllables drops significantly during learning (Fig. 1D, pink bars). Spike rate variability also dropped significantly during the same period (Fig. 1D, gray bars), along with other developmental changes in spiking statistics including the emergence of bimodality in the distribution of inter-spike intervals (Fig. S1A) and increased sparseness in spiking during song (Fig. S1B). These parallel previous findings from juvenile zebra finches [10] and from adult zebra finches and Bengalese finches [26–30]. While these observations highlight the gross changes in RA activity across development, they do not explain whether these changes reflect a developmental change in the underlying motor code.

To address this gap, we quantified whether the timescale of motor coding in RA changes during learning. As in our prior work [11, 29], for each song syllable we chose a time point after the onset of the syllable at which we measured the acoustic features of each rendition (see Song Acoustic Analysis). We then analyzed RA neurons’ spiking during the syllable’s “premotor window” (40 ms preceding the time at which a syllable’s acoustics were measured), during which RA is known to shape syllable acoustics.

### Millisecond-scale motor coding is present throughout learning

To examine the timescale of the vocal motor code, we quantified how well RA spiking predicts syllable acoustics when spike times in the premotor window are binned at progressively finer temporal resolutions (Fig. 2). To do so we used Mutual Information (MI), a model-independent measure of statistical dependence between two variables. For each syllable rendition, we quantified three acoustic features: pitch, amplitude, and spectral entropy. These syllable features vary from trial to trial under neural control and are refined during vocal learning [15, 29, 31, 32]. For each syllable, individual renditions were defined as “low” or “high” by a median split of that feature (see Mutual information estimation and entropy-based variability). Simultaneously, each syllable-paired trial of RA activity from the 40-ms premotor window was discretized into bins of width *dt* milliseconds, and each rendition was represented as a “word”—a vector of the counts of spikes across those bins (Fig. 2A). This set of spiking-behavior pairing yields a syllable-neuron “case,” where each case is the set of spike words emitted by an RA neuron and the paired set of binarized acoustic output of a given syllable.

**Fig. 2.**
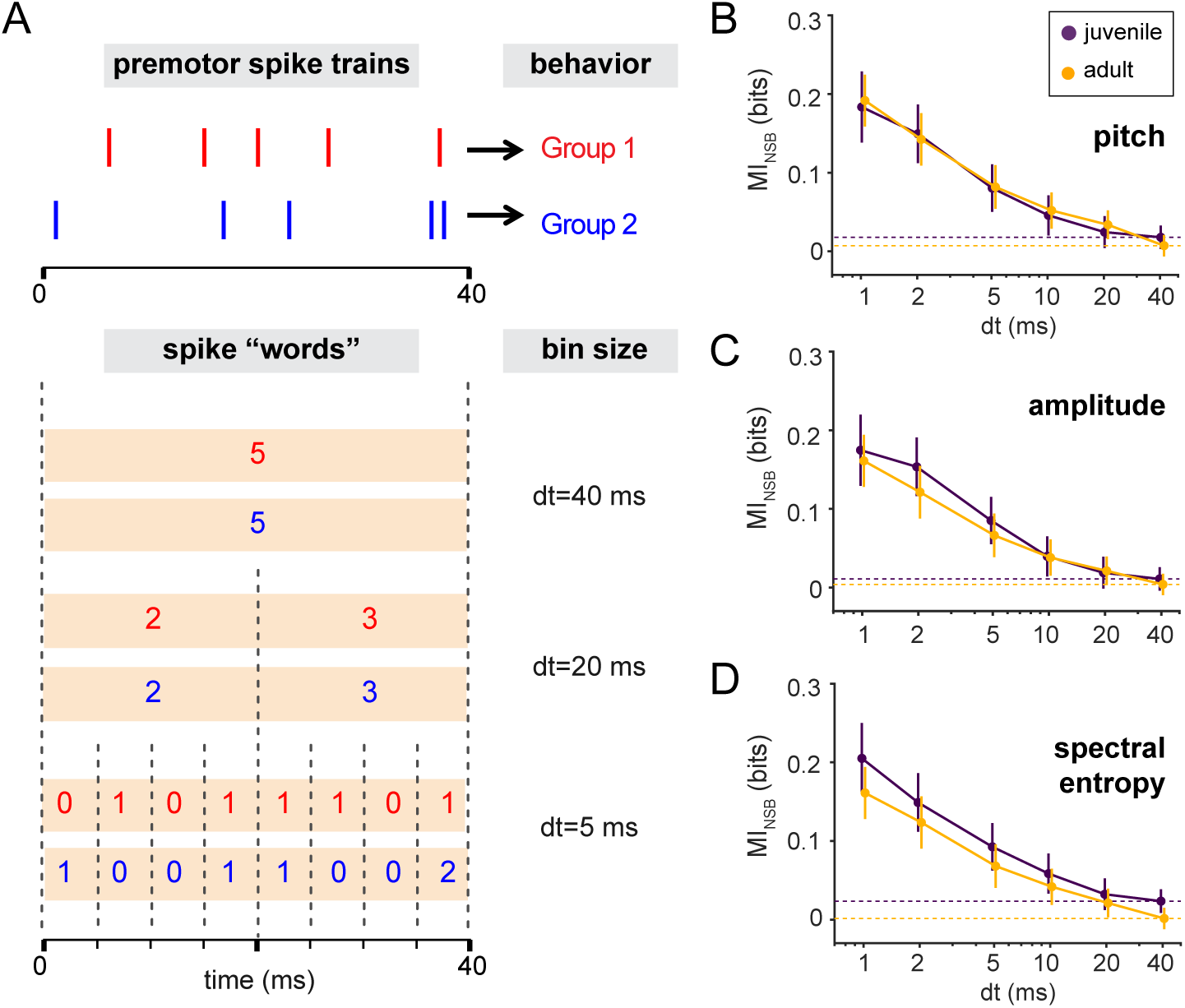
A precise timing code for vocal behavior control is observed in both juvenile and adult birds. **A)** Premotor spike trains (top) associated with distinct behavior groups (red vs. blue) are converted into spike “words” by binning spike counts at varying temporal resolutions (bin sizes, *dt*). At coarse resolutions (*dt* = 40, 20 ms), both spike trains are identical words (5; [2,3]), whereas a finer resolution (*dt* = 5 ms) reveals words of distinct temporal patterns. **B-D)** Mutual information (MI) between the spike words and acoustic features at different temporal resolutions *dt*, as estimated by the NSB estimator (see Mutual information estimation and entropy-based variability). The three panels display MI between spike words at different *dt* resolutions and three acoustic features: B) pitch, C) amplitude, and D) spectral entropy. Purple curves represent juveniles (51 neuron-syllable cases); gold curves represent adults (23 cases). Error bars represent one standard deviation (SD) of the information estimate (see Mutual information estimation and entropy-based variability). In all three acoustic parameters, MI increases as *dt* decreases (approaching 1 ms). For both juvenile and adults, MI at the coarsest temporal resolution (spike count in the entire premotor window, *dt* = 40 ms, dashed lines), are not statistically different from zero, indicating that variations in the spike rate do not encode variations in these acoustic features. Note that we consider two acoustic groups for each acoustic feature, thus the maximum possible MI is one bit.

Contrary to our hypothesis, in juvenile songbirds RA neurons encode vocal acoustics at millisecond precision rather than at coarser timescales. For each case, we computed MI between the spike words and the binary acoustic outcome and asked which timescale yields the highest MI. We found that MI was near-zero at larger bin sizes and increased only as we approached millisecond resolution. As shown in Figure 2B, MI between RA spiking and syllable pitch rose monotonically as bin size (*dt*) was reduced from tens of milliseconds to 1–2 ms in both juveniles and adults, with nearly identical values of MI at each *dt* across the two age groups. We found the same dependence on *dt* for syllable amplitude and spectral entropy (Fig. 2C–D), indicating that millisecond-scale timing is a general feature of RA’s motor code across acoustic dimensions. These results in juvenile birds contradict our hypothesis of a developmental change from rate-based to timing-based coding: for all acoustic parameters, bins coarser than ∼10 ms carried MI values not significantly different from 0 bits, and predictive information emerged progressively as *dt* approached the 1–2 ms regime, consistent with our prior findings in adult birds [11].

To determine whether the results shown in Figure 2 might reflect a gradual change in motor coding during the juvenile period, we subdivided cases into four age groups (65-69, 73-79, 80-88, *>*140 dph) and computed MI across *dt* for each age group for all acoustic variables we examined. We found no differences among age groups (Fig. S3). Notably, even the youngest age group exhibited high values of MI at *dt* = 1-2 ms and negligible MI for *dt* ≳ 10 ms, indicating that millisecond-scale control is present from early stages of sensorimotor learning and persists into adulthood. To test the robustness of our MI estimates, we considered two categories of potential sources of biases in our dataset [33]: sample size and data non-stationarities. We found that our MI estimations are robust to both potential sources of biases (Fig. S5-S8). Taken together, these results argue against a developmental shift from rate-based to timing-based control as an explanation for the coordinated reductions in vocal and spike-count variability (Fig. 1D). Instead, a millisecond-scale timing code is present early in sensorimotor learning, persisting *alongside* the decline in rate variability. We therefore predicted that rather than sharpening the temporal resolution of the motor code, learning might operate primarily by refining the repertoire of specific precise spike patterns.

### Evolution of RA spike pattern vocabularies across learning

To test this prediction, we asked what aspects of RA spiking change during learning (since the timescale of control is fixed; Fig. 2). Specifically, we assayed developmental changes in the size of each neuron’s “vocabulary” of precisely-timed spikes. We used the entropy of each neuron’s distribution of spiking patterns (at some particular *dt*) as a metric of vocabulary size. Specifically, we examined the entropy of neurons’ spiking vocabulary at two timescales: scalar spike counts (*dt* = 40 ms) and millisecond-resolved spike patterns (*dt* = 1ms). Unlike Fano Factor, which we used previously to quantify count dispersion and relies on Gaussian assumptions (see Spike count variability (Fano factor)), entropy is a model-free measurement of variability that can be applied to a distribution of spike counts or of 40-symbol binary spike words (see Entropy-based variability). This allowed us to compare variability in rate and in precise timing within a single framework.

At *dt* = 1 ms, spike pattern entropy did not decline with age; instead, it increased modestly with learning (Fig. 3A). This result is initially surprising, as learning is often associated with a reduction in variability. However, this reduction has been measured only at coarse timescales to date, whereas the statistics at finer temporal resolutions can evolve differently (see schematic in Fig. 4). To reconcile these observations —the increase in entropy at small *dt*, the reduction in variability of spike rates, and the stable timing precision of the motor code during learning —it is important note that entropy depends on the mean number of spikes in the premotor window. As more spikes are present, the number of ways they can be arranged into distinct patterns increases dramatically (e.g. a 40-ms spiking window binned at *dt* = 1ms yields 40 time bins each with a 0 or 1 spikes), which can increase the entropy (exemplified in Fig. 4). Consistent with this idea, we observed that the mean spike count within the premotor window did, in fact, increase with age (Fig. 3B) and that spike-pattern entropy increased with higher mean spike counts (Fig. 3C).

**Fig. 3.**
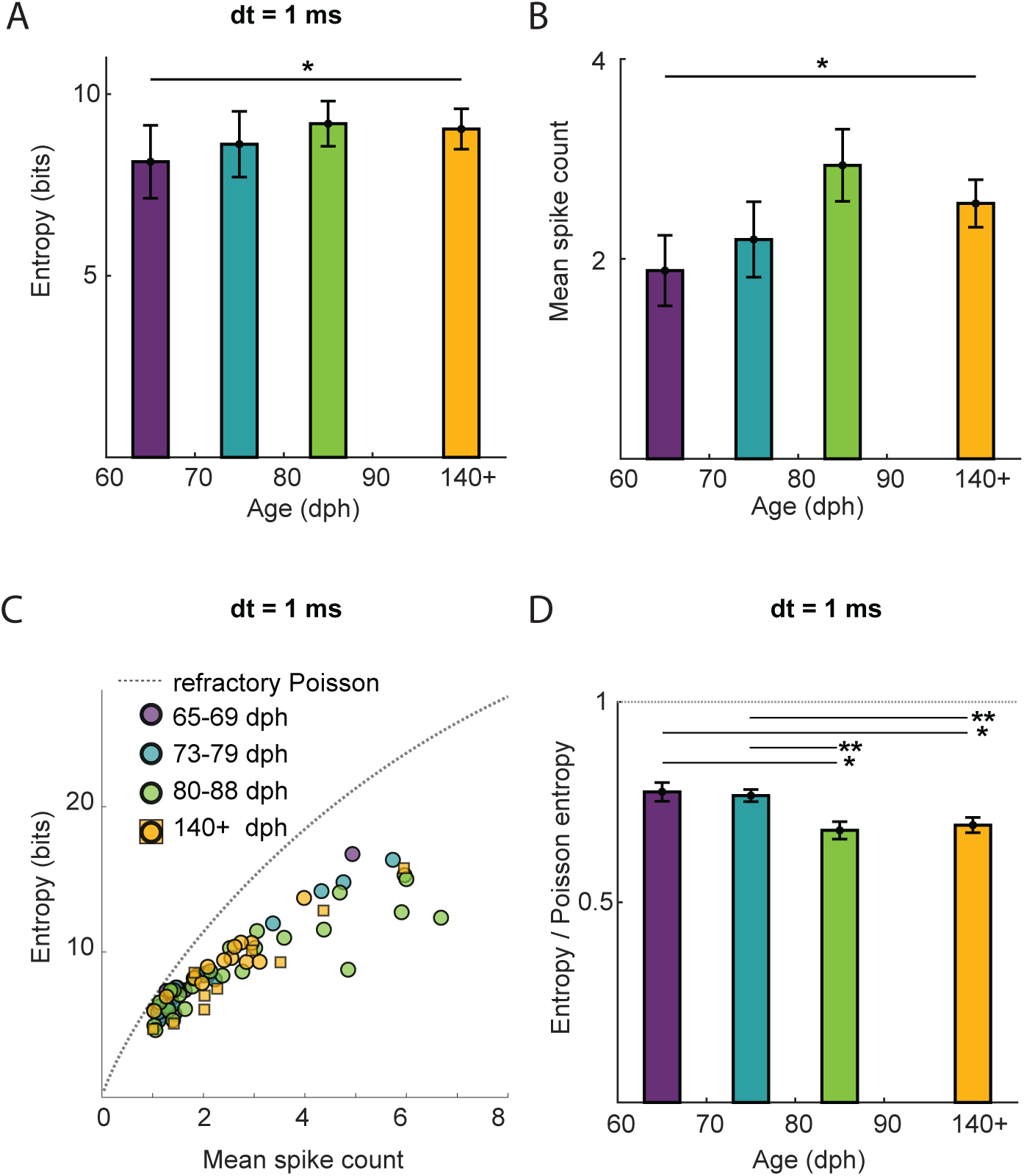
Refinement of millisecond-resolved spike patterns across learning. **A)** Raw 1-ms spike-word entropy increases across development, indicating an expansion of total pattern diversity. **B)** Mean spike count during the 40 ms-long premotor window increases across development. **C)** Relationship between mean spike count and entropy relative to a refractory Poisson reference model (dashed line). Entropy increases with firing rate but progressively deviates below the Poisson prediction in older birds, indicating the emergence of temporal structure. Circles represent longitudinally tracked birds; squares represent adult cases from previous data [11]. **D)** Normalized entropy decreases with age. This deviation from the rate-matched Poisson model indicates that spike patterns become increasingly selective and structured during sensorimotor learning, rather than simply reflecting changes in firing rate. Colors indicate age group: 65–69 dph (*n* = 10), 73–79 dph (*n* = 16), 80–88 dph (*n* = 25), and 140+ dph (*n* = 23). In **A**, **B**, and **D**, statistical significance was assessed via two-sample KS tests between age groups. Asterisks denote significance levels: * *p <* 0.05; ** *p <* 0.01. Error bars represent mean ± SEM.

**Fig. 4.**
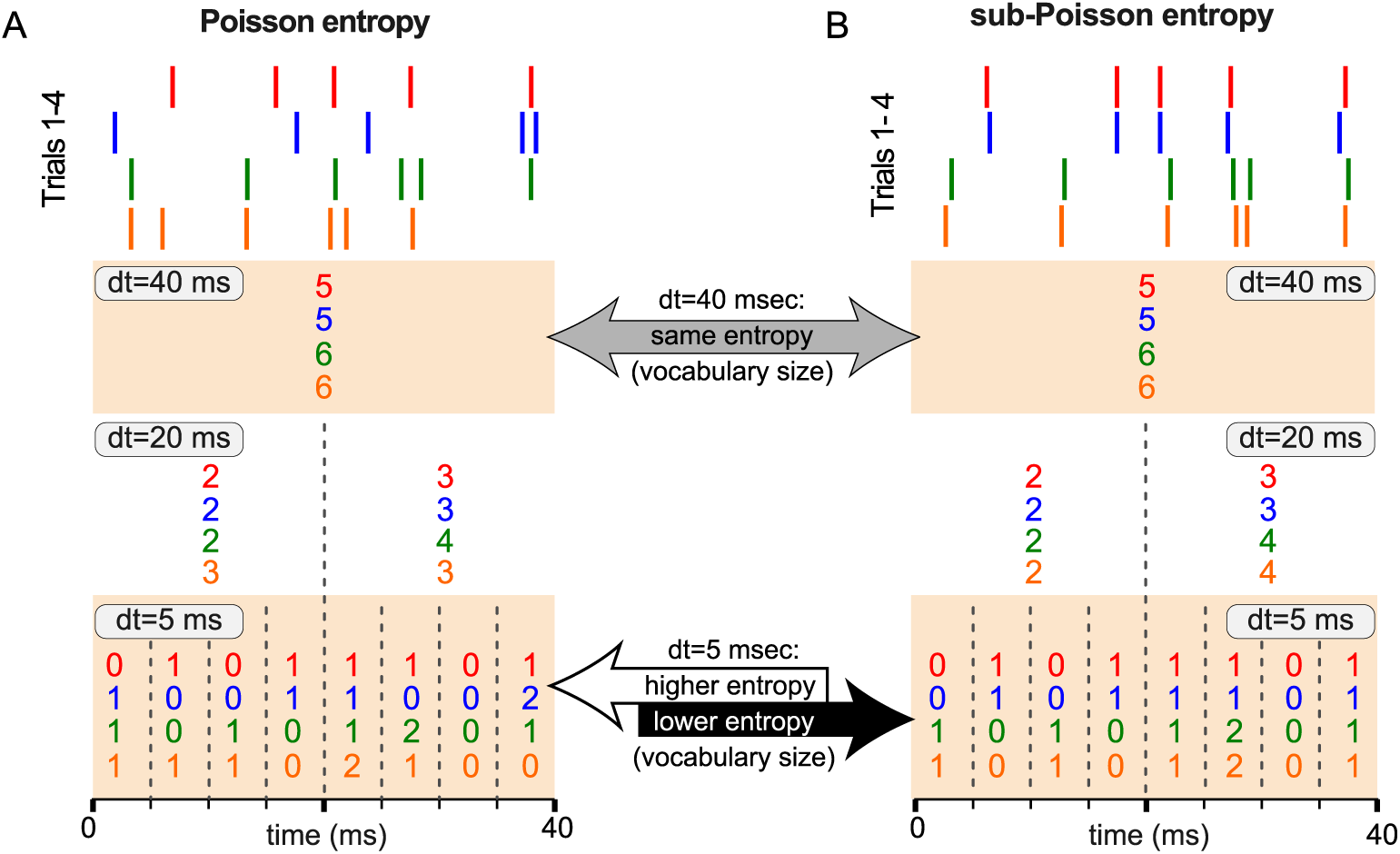
Temporal resolution reveals pattern-level structure beyond spike count. Raster lines show four example trials from a single RA neuron within a 40-ms premotor window. Colored numbers depict the spike count in each bin at resolution *dt* (coarse: *dt*=40 ms; intermediate: *dt*=20 ms; fine: *dt*=5 ms). **A) Poisson entropy:** Example trials generated by a refractory Poisson process. At *dt*=40 ms, all trials share the same total spike count and are therefore indistinguishable. As *dt* decreases, the same spikes can occupy different time bins across trials, yielding many distinct spike words (a larger vocabulary) and higher entropy. **B) sub-Poisson entropy:** Example trials matched to panel A in total spike count (and thus entropy) at *dt*=40 ms, but with more reproducible timing across trials at finer *dt*. This increased trial-to-trial alignment collapses the set of observed spike words, producing a smaller effective vocabulary and lower entropy at high temporal resolution than expected from a refractory Poisson process with the same rate.

To dissociate changes in spike pattern entropy from the increase in the overall firing rate, we normalized entropy for each case by the entropy of a rate-matched refractory Poisson process (see Methods). This normalization controls for mean rate and provides a benchmark for temporal structure (values near 1 indicate Poisson-like timing variability, whereas lower values reflect more temporally structured spiking). Since Poisson entropy is directly proportional to spike rate, this normalized metric is closely related to a bits per spike measure used in similar analyses [34]. A clear developmental signature emerged: normalized entropy decreases modestly but significantly with age (Fig. 3D). This indicates that relative to Poisson spiking, neurons’ vocabulary of precise spike timing patterns becomes increasingly selective as learning progresses. Considered alongside the MI results (Fig. 2), these entropy trends support a temporally resolved view of motor learning: rather than shifting control from coarser to more precise timescales, millisecond-scale precision is present throughout development, and sensorimotor practice shapes behavior by selecting and stabilizing a more structured and stereotyped repertoire of precise spike-timing patterns.

## Discussion

Acquiring a motor skill requires the nervous system to build a vocabulary of neural spike patterns that control behavior. By recording from single RA neurons across the sensorimotor learning period in Bengalese finches, we asked whether the temporal precision with which these spike patterns shape behavior (i.e., the motor code) changes during learning. We found that the motor code for birdsong is already millisecond-precise at the youngest ages we can analyze, leading us to speculate that learning proceeds mainly by selecting which precise timing patterns are used, rather than by refining the timescale of the code itself.

Our results bear on a central question in the field of motor control. Most prior work on motor coding during skill acquisition has emphasized changes in rates, SNR, or ensemble covariance [9, 35, 36])—largely assuming motor coding on coarse (tens to hundreds of milliseconds) timescales from adult animals. This left open the question of whether fine temporal control emerges during learning. Our MI analyses demonstrate that the neural–acoustic correlation in juvenile birds is negligible at coarse resolutions but is strongly predictive of behavior when spikes are examined at 1–2 ms resolution (Fig. 2B-D, purple curves). The millisecond-scale precision previously shown to underlie skilled behavior in adult songbirds [11] therefore also underlies vocal behavior in juvenile birds who are still learning their songs (Fig. 2).

These findings falsify our initial hypothesis that developmental changes in spiking reflect a gradual sharpening of the vocal organ’s biomechanical tuning. Instead, they suggest that learning proceeds by exploiting a pre-existing, millisecond-precise substrate through the selection and stabilization of specific spike patterns.

Future studies could examine whether vocal learning in adult animals (e.g. sensorimotor adaptation or recovery from neural injury) exhibits similar or distinct changes in the timescale of the motor code (Fig. 2) or the entropy of the spiking vocabulary (Fig. 3). Although millisecond-scale precision has not been systematically tested during learning in adult songbirds, our results imply that adult vocal learning likewise relies on millisecond-precise control—implemented through pattern selection rather than shifts in coding timescale. Beyond songbirds, future work in other species and behaviors might apply this approach to ask whether other forms of developmental learning are (or are not) accompanied by changes in the intrinsic timescale of the motor code and/or the vocabulary of spike patterns.

The force-producing properties of muscle fibers determine how neural spike trains shape behavior. Notably, syringeal muscles exhibit “superfast” twitch dynamics before the onset of sensorimotor learning, indicating that millisecond-scale spike timing can indeed shape force production early in development [18, 20]. Moreover, passive (non-muscular) sound-production properties of the syrinx show no developmental change in zebra finches [21]. Therefore, although we cannot rule out the possibility that dynamics of muscle force production might still be changing during the ages at which we have recorded RA neurons, these findings suggest that the biomechanics of the syrinx are stable during the learning period we studied. However, we do not yet know if this is the case during subsong, the earliest, “babbling” phase of vocal development. Future experiments recording from RA during this time—before identifiable syllables emerge—might clarify whether the motor code is fixed from the very first vocalization or if it develops alongside the bird’s earliest attempts to sing.

These findings help reconcile the long-standing observation that acoustic variability and spike-count variability co-reduce during learning (Fig. 1D, Fig S2). Prior studies have only examined spiking on coarse timescales and, in doing so, implicitly treated rate variability as the primary biologically relevant signal. At coarse resolutions, many distinct millisecond-scale spike patterns collapse onto the same spike-count representation, whereas at fine resolutions those differences expand the effective pattern vocabulary (Fig. 4). In contrast, our approach does not impose this assumption. Instead, it reveals that behavior is best predicted by millisecond-scale spike patterns rather than by slower rate fluctuations (Fig. 2).

Crucially, the reduction in the Poisson-normalized entropy we observed indicates that learning involves a contraction of the effective spike-pattern repertoire relative to chance. As illustrated schematically (Fig. 4), matching spike-count statistics can coexist with large differences in pattern-level entropy, and sub-Poisson entropy at fine *dt* reflects increased trial-to-trial reproducibility of specific timing motifs. This suggests that the decline in neural variability is not merely a reduction in noise, but rather reflects the active selection of specific, structured timing motifs from a broader range of possibilities. Future studies might test this prediction by “writing in” precisely-timed spike patterns, for example with holographic optical stimulation [37–39]. Such an approach would allow for the direct tests of how specific timing shifts influence vocal output, thereby confirming the causal necessity of the precise motor code we have described.

The observed reshaping of the spiking vocabulary (Fig. 1C-D; Fig. 3; Fig. S1,S2,S4) may be attributed to developmental changes in the synaptic architecture rather than the fundamental physiology of RA neurons. While the intrinsic excitability of RA neurons changes significantly early in development, these shifts are largely completed by 50 dph [40, 41]—prior to the age at which individual syllables can be reliably discriminated. After 50 dph, instantaneous firing frequency responses and spike rate adaptation remain stable, with only minor changes in spike threshold and waveform [41]. Consequently, the RA neurons we recorded during the sensorimotor learning period likely maintain a consistent ability to respond to external input. This stability suggests that the emergence of precise patterns is driven by changes in the drive from two major inputs (Fig. 1B): LMAN, which guides exploration via the anterior forebrain pathway [10, 22, 32, 42], and HVC, which provides timing signals via the motor pathway [28, 43, 44].

Developmental plasticity in these upstream inputs offers a plausible substrate for the pattern selection we observe. In the anterior forebrain pathway, LMAN spiking becomes more song-locked and burst-prone over development [8, 40, 44, 45]. Simultaneously, in the motor pathway, HVC→RA connectivity is refined through strengthening and pruning [40, 46]. Modeling and circuit evidence suggest that this reorganization reduces rate variability and enhances temporal reliability [40], while the recruitment of HVC interneurons tightens the temporal structure of the drive sent to RA [47–50]. Locally, the selective pruning of inhibitory circuitry within RA suggests that intra-RA inhibition may act to “carve out” specific allowable patterns from broad excitatory drive [51]. Together, these convergent changes in LMAN, HVC, and local inhibition might select and stabilize specific millisecond-precise patterns without altering the underlying temporal resolution of the motor code.

We note several methodological limitations to our study. First, we did not track the same neuron across days; while our cross-sectional approach reveals population-level shifts, the most direct test of pattern selection within individual neurons will require long-term, stable single-cell recordings spanning the entire learning trajectory. Moreover, although we used stringent controls to limit estimator bias (See Supplementary Information, Fig. S5-8), MI estimation at millisecond resolution is inherently data-hungry; larger datasets would be needed to help narrow confidence intervals on age-by-timescale effects. Recent advances in neural-network-based MI estimation techniques may also prove valuable for future analyses [52].

In summary, our results demonstrate that the temporal precision of the songbird motor code is present early, suggesting that learning proceeds chiefly by selecting which precise multi-spike patterns are expressed. Song learning thus may be a progressive curation of a millisecond-resolved spike-pattern vocabulary. This framework parsimoniously accounts for the co-reduction in spike-count and acoustic variability, the millisecond-limited information curves, and the rate-matched contraction of spike-word entropy. More broadly, it advances a falsifiable model for vocal control and possibly other complex motor skills: that the timescale of control is fixed, and performance improves by refining the repertoire used on that timescale.

## Methods

### Subjects

Male Bengalese finches (*Lonchura striata var. Domestica*, a songbird species of which only males learn to sing) were bred in our colony and housed with their parents until 60 days of age. After electrode implantation, birds were isolated and housed individually in sound-attenuating chambers with food and water provided *ad libitum*. All recordings are from undirected song (i.e., no female was present). Recordings from five juvenile birds (60–90 dph) were collected using a previously described experimental protocol [29]. Although sensorimotor learning in Bengalese finches typically begin ≈ 40 dph [53, 54], we chose 60 dph as the earliest age for our analysis for two reasons. First, song syllables from days earlier than 60 dph are acoustically difficult to quantify and categorize into distinct syllable identities, constraining our ability to quantify within-syllable acoustic variation. Second, the surgical challenges of implanting younger Bengalese finches include their fragile, immature skull: attempting to secure a microdrive on their skulls at younger ages would compromise our ability to hold single neurons for longer periods as well as the sturdiness of the implant through the weeks-long chronic recordings. Recordings from adult animals (*>* 140 dph) were obtained from one of the birds that was recorded as a juvenile and matured into adulthood, and from three adult birds that were previously obtained as part of a separate analysis [29]. Procedures were performed in accordance with established animal care protocols approved by the Emory University Institutional Animal Care and Use Committee.

### Song Acoustic Analysis

Birdsong was chronically recorded with a microphone placed inside a sound-attenuated housing chamber, along with neural recordings. To quantify the acoustics of song syllables, we first detected the presence and identity of song syllables from the audio recordings. Syllable onset and offset times were determined on the basis of amplitude threshold crossings after smoothing the acoustic waveform with a square filter of width 2 ms. This method allowed for the determination of syllable onset times with millisecond-scale precision [11].

### Syllable Acoustic Quantification

The identities of song syllables were determined by visual examination of spectrogram renderings of song behavior. For each syllable identity, we selected a specific time point (relative to the detected syllable onset) when the vocal acoustic features (e.g. pitch during a sustained note) were most stable and well-defined. This syllable-specific time point of acoustic quantification was fixed relative to the syllable onset (the latency between syllable onset and time of quantification was constant across all iterations of a given syllable type).

We analyzed the acoustic power spectrum at the specified time for each iteration of a song syllable to quantify the fundamental frequency (which we refer to as “pitch”), amplitude, and spectral entropy. We chose to quantify these three acoustic features since they capture a large percentage of acoustic variability in Bengalese finch song and are the features that are refined during song learning [25, 29].

### Electrophysiological recordings

Birds were anesthetized (induction with 3% isoflurane, maintained with 1.5–2% isoflurane) and a lightweight 16-microelectrode bundle microdrive (Innovative Neurophysiology) was stereotactically positioned above RA nucleus in one hemisphere (2 implants over right RA, 1 over left RA) and secured to the skull with dental cement. After birds recovered from surgery and singing resumed (within 1–3 days), electrodes were advanced through RA using a miniaturized microdrive which recorded extra-cellular voltage traces during and between bouts of singing. RA recording sites were confirmed by the presence of characteristic changes in activity associated with the production of song and calls and by *post hoc* histological confirmation of electrode tracks passing through the RA nucleus.

### RA Unit Isolation and Inclusion Criteria

To isolate spiking activity from individual units, we used a previously described spike sorting pipeline [29] that yields a scalar isolation measure for quantitative inclusion. Briefly, we set a voltage threshold to detect both spike and noise waveforms during singing-related activity, performed principal components analysis (PCA) on detected waveforms, and projected each waveform onto the first two principal components. We then assigned waveforms to clusters using an automated nearest-neighbor clustering algorithm (kmeans.m in MATLAB, The MathWorks, MA) applied to the 2-D feature space; in most cases, two clusters were selected (a “spike” and a “noise” cluster).

We quantified unit isolation by measuring overlap between spike and noise clusters. Specifically, we fit a 2-D Gaussian to each cluster to estimate its mean and covariance, generated 10,000 synthetic points from each Gaussian, re-applied the nearest-neighbor classifier to the synthetic points, and defined “isolation error” as the fraction of synthetic points misclassified by the algorithm. Units with isolation error *<* 0.01 were classified as single units.

To benchmark the stringency of this overlap-based criterion against other commonly used cluster-quality metrics, we additionally computed the *L-ratio* and *isolation distance*, which quantify separability using Mahalanobis distances in feature space [55, 56]. Consistent with these alternative measures, we evaluated L-ratio for the sub-set of recordings closest to our inclusion cutoff (i.e., the worst-isolated recordings among those that nonetheless passed the *<* 1% overlap criterion) and found that these recordings fall within or below ranges previously considered well isolated. Moreover, within this “worst” subset by our overlap metric, a majority of cases still exhibited low L-ratio values indicative of good isolation, indicating that our overlap-based inclusion criterion is conservative relative to these commonly used metrics.

We next applied physiological and cell-type inclusion criteria. We examined the inter-spike interval (ISI) distribution of each isolated unit and included only units with *<* 1% of ISIs shorter than 1 ms. Based on prior characterizations of RA cell-type-specific spike waveform shape and response properties [29], recordings were classified as putative excitatory projection neurons or interneurons; *>* 97% of isolated units were classified as putative projection neurons. Only RA projection neurons were included in analyses.

Across 8 Bengalese finches, our dataset comprised 67 single units recorded across ages spanning 62 dph to *>* 140 dph. Adult RA neurons were obtained from three birds from a previous study [29] at ages *>* 140 dph (pu26y2: 6 neurons; pu24w39: 11 neurons; pu44w52: 17 neurons) as well as a juvenile bird that matured into adulthood (Bird2: 2 neurons). Longitudinal recordings during sensorimotor learning were obtained from Bird1 (16 neurons) spanning 26 days (62–88 dph) and Bird2 (10 neurons) spanning 77 days (78 dph–155 dph). Additional juvenile birds contributed shorter spans: Bird3 (2 neurons) spanning 2 days (79–81 dph), Bird4 (4 neurons) spanning 3 days (85–88 dph), and Bird5 (1 neuron) on a single day (76 dph).

### Spike train visualization and analysis

To visualize the consistency of neural activity across the motif, spike trains were aligned to the onset of the first syllable. To account for natural temporal variability in song production, we applied a piecewise linear time-warping algorithm [29]. This process partitioned the motif into segments based on acoustic landmarks (syllable onsets and offsets). We calculated a linear scaling factor for each syllable and inter-syllable interval to map its duration to the corresponding segment of a mean duration reference motif. This alignment ensured that both the target syllable and downstream acoustic landmarks were visually aligned across trials. Crucially, this warping was performed strictly for visualization (Fig. 1C); all quantitative analyses were conducted on unwarped spike times relative to specific syllable onsets.

#### Premotor window definition

For all analyses of neural activity [11, 19], we defined a premotor window of length *T* = 40 ms, relative to the time of quantification for acoustic features. This window captures the relevant motor command generation for the associated syllable.

### Spike count variability (Fano factor)

To quantify trial-to-trial variability at the coarse timescale of the full premotor window (*T*), we calculated the Fano factor [57]:

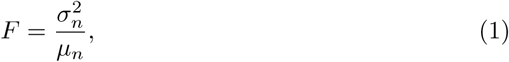

where *n* is the total spike count in window *T*, *σ*^2^*_n_* is the variance, and *µ_n_* is the mean count across trials. The Fano factor compares the observed variability to that of a homogeneous Poisson process (*F* = 1). Note that this spike-count metric does not capture refractory effects or the temporal structure of spiking at finer timescales.

### Mutual information estimation and entropy-based variability

To determine the temporal scale at which premotor spiking predicts acoustic variation, we estimated the mutual information (MI) [58] between discretized measures of pre-motor spiking activity and behavioral groups of syllable acoustics (Fig. 2A) [11, 19]. Because MI is computed from (conditional) entropies of spike “words,” the same entropy estimates also provide a natural, time-resolution-dependent measure of neural variability. Accordingly, after identifying the millisecond timescale as behaviorally predictive (Fig. 2B-D), we used the (unconditional) spike-word entropy at the relevant *dt* as a complementary measure of neural variability that captures temporal pattern structure beyond spike-count statistics (Fig. 3).

#### Discretization

Within the premotor window *T*, we discretized the spike train into bins of duration *dt*, with 1 ≤ *dt* ≤ 40 ms (Fig. 2A). Given the refractory period of RA neurons (*τ*_ref_ ≈ 2 ms) [41, 51], we expect no more than one spike per 1 ms bin; however, larger *dt* bins can contain multiple spikes. Therefore, we digitized the spike times such that the value in each bin represented the spike count. This resulted in a “spike word” *R* of length *T/dt*. To examine spike trains across temporal resolutions, we repeated this discretization for a range of timescales *dt* = {1, 2, 5, 10, 20, 40} ms.

#### Behavioral groups (G)

In this study, the behavioral variable *G* was defined based on the acoustic features of individual syllables. To quantify the sensitivity of premotor spiking to acoustic variation, we generated binary behavioral groups: for a given syllable, individual renditions were labeled “low” or “high” based on a median split of the distribution of the acoustic feature being analyzed.

#### Mutual Information calculation

Using *N*_trials_ spike words *R* of size *T/dt*, together with the *N*_trials_ acoustic group indices *G* for each acoustic feature, one can then estimate MI between the spike train and the motor output through a difference in two entropies [11]:

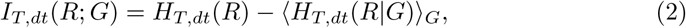

where ⟨*. . .* ⟩*_G_* represents an average over the behavioral group *G*. MI, Eq. (2), quantifies the reduction in uncertainty of a random variable *R*, as measured by the entropy *H*(*R*), due to the knowledge of another random variable *G*, as quantified by the *conditional* entropy *H*(*R*|*G*) [59].

Given a probability distribution *p*(*R*), the entropy *H*(*R*) is defined as

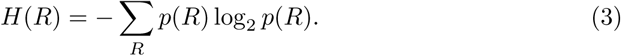

Obtaining an accurate estimate of the entropy from finite datasets is a notoriously difficult problem [33], as it requires an estimation of the probability distribution *p*(*R*) from a finite number of samples. In the asymptotic sampling regime, where each response *R* has been sampled multiple times, the “naive” or “maximum likelihood” (plug-in) estimate can be used, which replaces probabilities by empirically observed word frequencies *p*(*R*) ≈ *f* (*R*) = *n_R_/N*_trials_ (where *n_R_*is the number of times a specific response *R* was observed, and *N*_trials_ is the total number of trials) in the definition of the entropy, Eq. (3). Such an estimate of the entropy is *downwardly biased* for finite datasets (i.e., it systematically underestimates the true entropy), with the bias converging to zero as *N*_trials_ → ∞ [33]. The convergence is slow, scaling approximately as ∼ |*R*|*/N*_trials_, where |*R*| is the cardinality of the variable whose entropy is being analyzed.

Because MI is computed as a difference of entropies, finite-sample bias in the individual estimates *H*(*R*) and *H*(*R*|*G*) can propagate into *I_T,dt_*(*R*; *G*) in a nontrivial way: although both entropies are underestimated by the plug-in estimator, their biases need not cancel, and naive MI estimates are therefore not guaranteed to be unbiased at finite *N*_trials_ [33].

The cardinality of possible responses depends exponentially on the size of the time window *T* and the time binning resolution *dt*, and can, in principle, be very large, precluding the use of the naive estimator. Indeed, we estimate the cardinality |*R*|_est_ of the observed spike trains using the standard estimate for the number of “typical” sequences when drawing *n* independent and identically distributed discrete random variables (known as the “asymptotic equipartition property”) [59]:

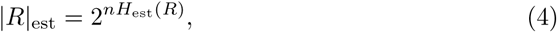

where *n* = *T/dt* is the number of bins used to discretize the spike trains, and we take the entropy estimate *H*_est_(*R*) to be the entropy of a Poisson distributed counts for the number of spikes in each bin with mean 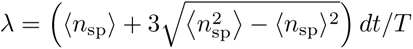, where *n*_sp_ is the total number of spikes observed during the premotor window *T* in each trial, and ⟨*. . .* ⟩ represents an average over trials. At the finest discretization considered here, *dt* = 1 ms, and for a typical average spike count, we find the number of possible neural words is on the order of |*R*|_est_ ∼ 10^10^. As this cardinality is much higher than the number of samples available per behavioral group, we necessarily operate in a severely undersampled regime. Following [11], we therefore required a minimum of *N*_trials_ ≳ 200 trials per group to ensure sufficient coincidences for reliable entropy estimation.

#### NSB entropy estimation and long-tail partitioning

To estimate *H*(*R*) and *H*(*R*|*G*) in this undersampled regime, we used the Nemenman–Shafee–Bialek (NSB) entropy estimator [23, 60, 61]. NSB is a Bayesian approach that infers entropy from the observed histogram of word counts and, in practice, can strongly reduce finite-sample bias relative to the plug-in estimator in undersampled settings by leveraging the number of repeated observations (“coincidences”) in the dataset [11, 23]. We verified that the coincidence criterion required for reliable NSB estimation was satisfied for all syllables and acoustic groups included in the analysis; specifically, we required a minimum of 200 trials per syllable, and additionally restricted analyses to syllables with mean spike count *>* 1 to ensure the unit was “active” [29]. A total of 74 neuron-syllable cases across 5 birds and ages from 65 dph to *>* 140+ dph satisfied these criteria.

The NSB approach to estimating the entropy relies on having a fast (e.g., exponential) decay of the rank-ordered distribution of words as a function of the rank [23, 61] (see also [62]). In particular, entropy estimates for rank-ordered distributions of words with long (e.g., power law) tails tend to underestimate the true entropy [63, 64]. To circumvent this problem, we make use of an exact relation for the additivity of entropy for a mixture of two disjoint partitions of data (i.e., containing no overlapping words), see Supplementary Information for a derivation of the formula for the additivity of entropy and the corresponding error bars on the total estimate. Thus, similarly to [11, 64], we partitioned trials into the most common word vs. all other words in cases (i.e., for a given syllable on a given day) where the most common spike word in the distribution appeared in more than 2% of the recorded trials. For juvenile birds, the most common word is in all cases the “silence” word with no spikes.

#### Bias controls

Because *I_T,dt_*(*R*; *G*) is computed as a difference of entropies, we performed additional controls to confirm that any potential finite-sample bias in *H*(*R*) and *H*(*R*|*G*) does not appreciably bias MI estimates. To do so, we verified that: 1) spiking statistics were approximately stationary across trials of each neuron’s recorded sample set, so that different spike words can be assumed to be independent samples of the underlying word distribution, and 2) entropy estimates do not suffer from undersampling bias. We give a short description of these two tests below, following [11], while the detailed results of these tests are provided in the Supplementary Information.

We tested for nonstationarities and correlations across trials of spiking data for every syllable/neuron case by comparing the difference in entropy estimates between the first and second half of all trials with the difference in entropy estimates between even and odd trials. The latter provides a reference in which any contribution from long time correlation effects should be canceled out. Cases in which half-split differences systematically exceeded even-odd differences would indicate potential nonstationarity; we did not observe such systematic deviations (Supplementary Information).

To address potential finite sample bias, we verified that individual conditional and unconditional entropy estimates showed little variations when computed from smaller data fractions than our complete dataset [11, 34, 64]. More specifically, when estimating the entropy for a case with *N* number of trials recorded, we also estimated the entropy *H*(*α*) for *αN*, with *α <* 1, randomly selected trials and averaged over 10 realizations of this subsampling to yield the subsampled-average entropy estimate ⟨*H*(*α*)⟩. By observing the finite data size scaling behavior of ⟨*H*(*α*)⟩ as a function of the inverse data fraction 1*/α* → 1, we could confirm that a large fraction of our entropy estimates were in an asymptotic regime of sufficient data size.

#### Averaging across cases

To obtain the MI curves for each age range in Fig. 2, we averaged the MI estimate in Eq. (2), first over all cases recorded in each day, and then over the different recorded days in each age range. The averaging in both cases was performed by inverse-variance weighting, i.e., by weighting the entropy contribution of each case by the inverse of the posterior variance of its estimate, as in [11]. The error bar on the MI average for each age range in Fig. 2 represents the standard deviation of the mean, and was similarly calculated by averaging the individual posterior variance of each estimate, first for all cases in each day, and then over the different recorded days in each age range, again weighted by inverse posterior variance. To further confirm the absence of undersampling bias in our data, we repeated the data fraction analysis described above on the final averaged MI estimates. We observe that our main finding from Fig. 2—namely that MI between spike timing and behavioral variation within a given syllable is higher at finer resolution, *dt* = 1 ms—is preserved when we subsample our dataset down to 50% of its full size (Fig. S5).

#### Entropy-based variability

To quantify neural variability in the spike-word space at each timescale *dt*, we compared the (unconditional) entropy estimated from spiking data within *T* to the entropy expected under a refractory Poisson reference process with refractory period *τ*. When *dt* = *T*, spike words reduce to spike counts, and the entropy of spike words is equivalent to the entropy of counts; in this case, the reference entropy can be computed exactly from the analytical count distribution for a Poisson process with a refractory period [65]. In the fine-resolution limit (*dt* = 1 ms ≪ *T*), we used a previously derived approximation for the entropy of spike words under a refractory Poisson model [63]. The refractory period used in the reference model was chosen to be compatible with reported refractory periods of RA neurons [51].

## Supplementary Information

This article has an accompanying Supplementary Information file.

## Acknowledgements

We thank Daphne Zhu and Margarita Sison for their help with syllable identification and histology. This work was supported in part by the National Science Foundation (NSF CRCNS Grant No. 1822677 to I.M.N. and S.J.S), the National Institutes of Health (Grants R01-NS099375 to S.J.S., U24-NS126936 to S.J.S., R01-NS084844 to S.J.S., 5R01NS084844 - 07 to S.J.S. and L.M.P., and F99-NS141391-01 to L.M.P.), the Howard Hughes Medical Institute (HHMI GT15944 to S.J.S. and L.M.P.), the Robert W. Woodruff Foundation (Woodruff Fellowship to L.M.P.), and the Simons Foundation as part of the Simons-Emory International Consortium on Motor Control.

## Author contributions

L.M.P., I.M.N., S.J.S. and M.P. designed the research project. L.M.P., I.M.N., S.J.S. and M.P. developed the theory. I.M.N. and S.J.S. managed the research project. L.M.P., A.V. and M.P. performed experiments and data analyses. L.M.P., I.M.N., S.J.S. and M.P. wrote the manuscript.

## Data, Materials, and Software Availability

Processed data (neural spiking data from isolated neurons and syllable acoustic measurements) and analysis code will be deposited in Github. Due to size reasons (several hundreds of gigabyte of data), the raw audio and neural recordings used in this paper will be shared by S.J.S. upon request. Further information and requests for resources and reagents should be directed to S.J.S. Part of the adult Bengalese finch data used in the present study were used for previous published work, where we demonstrated millisecond-scale motor coding in adult birds [11]).

## Supplementary Information

### Emergence of bursting activity during song acquisition in Bengalese finches

We found that late in the sensorimotor learning period (70–79 and 80–88 dph, blue and green traces in Fig. S1A) instantaneous firing rates (defined as the inverse of the interspike interval) show a bimodal distribution corresponding to high rates during bursts and low rates during inter-burst intervals. Early in sensorimotor learning, RA neurons produce a roughly exponential distribution of interspike intervals (ISIs) (62–69 dph, purple trace in Fig. S1A, inset).

To further characterize the reshaping of RA activity across song learning in Bengalese finches, we computed the sparseness index of firing patterns [1, 2], defined as 1 when the spiking is restricted to a single time bin during a song motif and 0 if spikes are evenly distributed across the motif (Fig. S1B). We selected one to three motifs from each bird’s song, with each motif representing a different sequence comprised of three song syllables. We found that the sparseness index increased during development from 0.069 ± 0.017 (*n* = 6) at 62-69 dph to 0.27 ± 0.09 in adult birds (140+ dph; *n* = 3). These results show that, as previously observed in zebra finches, during song learning juvenile Bengalese finches exhibit a transition in activity from exponentially-distributed ISIs in RA neurons – typical of a Poisson spiking model – with a low sparsity in the early stages of learning to a bimodal firing rate distribution and sparse activity in later stages, as RA activity becomes dominated by brief, high-frequency bursts separated by epochs of silence [3–5].

**Fig. S1.**
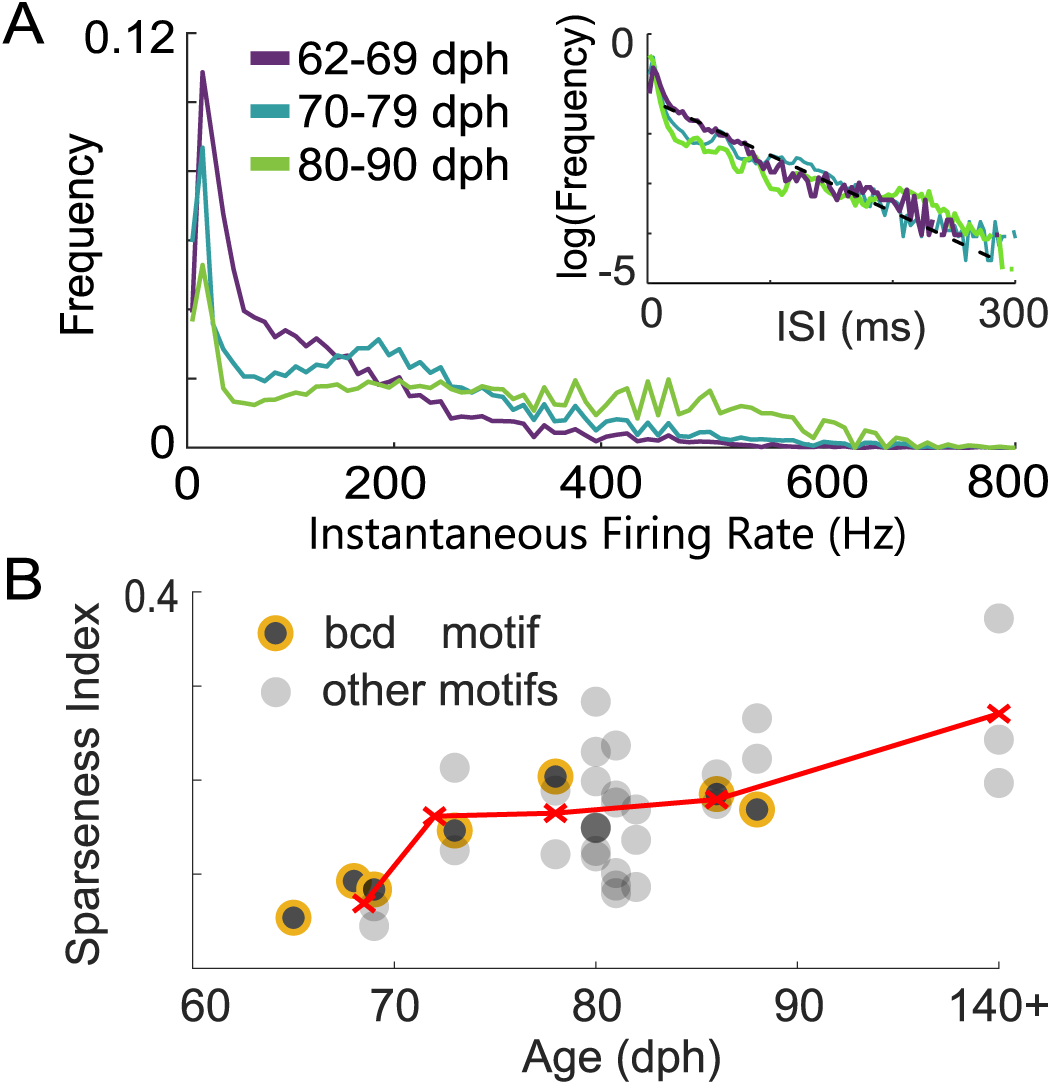
Transformation of gross RA spiking statistics in juvenile Bengalese finches across sensorimotor learning. **A)** Distributions of instantaneous firing rates (inverse of inter-spike interval, 1*/ISI*) are shown for three developmental timepoints: 62-69 dph (purple), 70-79 dph (teal), and 80-90 dph (green). Activity shifts from an approximately exponential, unimodal distribution in early learning to a bimodal distribution in late learning, reflecting the emergence of high-frequency bursting (*>*50 Hz, [5]). *Inset:* Log-frequency rendering of the ISI distribution show that early firing (purple) aligns with an exponential fit (black dashed line), consistent with a Poisson process, while later dates diverge significantly. **B)** Sparseness index (SI) [1, 2] of RA spiking increases across learning. Black circles represent the average SI of an RA neuron recorded across iterations of a motif. Circles outlined in orange represent SI of neurons recorded during the “bcd” motif shown in Fig. 1C. Red ‘x’ markers represent the average for different age groups from panel A.

### Spike rate variability and acoustic variability decrease simultaneously during song learning

Across all syllables and ages, the pitch distributions of juvenile syllables (62–90 dph) spanned a broad range of variability, including highly variable syllables (CV *>* 0.1), whereas pitch variability was substantially smaller in adults (140+ dph; Fig. S2B), consistent with prior work in adult Bengalese finches [5–7] and zebra finches [8]. We observed a parallel developmental trend in premotor spike count variability in RA: juvenile cases exhibited widely varying Fano factors, while adult syllable-days clustered at lower values (Fig. S2C). Importantly, variability in pitch and variability in spike count tended to co-vary at the level of individual syllables across learning. Because we cannot reliably track the same neuron across days, we quantify “syllable-day” measurements (i.e., the neuron(s) recorded for a given syllable on a given day), and we find that syllables that become more acoustically stereotyped across development also tend to be accompanied by reduced premotor spike-count variability in the RA recordings obtained for that syllable over learning. This positive relationship between pitch CV and spike-count Fano factor is exemplified by syllable ‘c’: from 69 dph to 88 dph (red stars in Fig. S2B–C), both pitch variability and spike-count variability decrease together, illustrating the broader co-reduction trend we observed across many syllables.

**Fig. S2.**
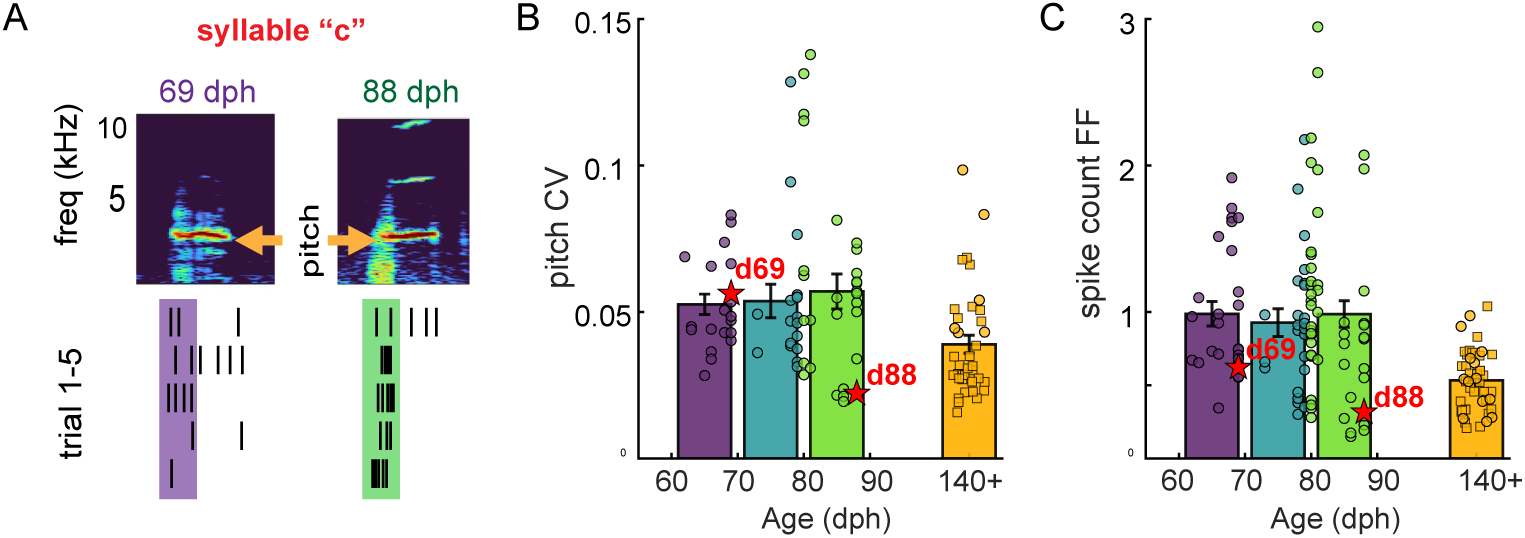
Concomitant reduction in pitch and spike count variability across learning. **A)** Representative spectrograms of syllable ‘c’ during early learning (69 dph) and late learning (88 dph) reveal changes in acoustic structure, which we quantify across learning (e. g., as changes in the fundamental frequency, which we refer to here as “pitch”). Similarly, changes in RA spiking activity during the 40 ms premotor window (shaded area) are quantified across learning. **B)** Pitch CV and **C)** spike count Fano factor for all analyzed syllables are depicted by individual circles. Values for the previously reported data [5, 9] are shown as square symbols. The error bars show the mean *±* SEM for each age group. Syllable ‘c’ at 69 dph and 88 dph is depicted with red stars.

### Entropy of RA spiking across learning

When spike patterns are analyzed at the scale of the whole premotor window (*dt* = 40 ms; i.e., spike counts in the entire premotor window), the entropy of spike-count “words” in both juveniles (65–88 dph) and adults (*>*120 dph) increases with mean spike count and tracks the prediction of a refractory Poisson process relatively closely (Fig. S4A). Consistent with reduced spike-count variability in adults, adult cases (*>*140 dph) tend to occupy lower-entropy values than juvenile cases at comparable mean spike count (orange symbols; Fig. S4A), and while a subset of juvenile cases lies above the refractory Poisson expectation, we did not observe any adult syllables with similarly high-entropy, “super-Poisson” spike-count behavior (Fig. S4A). In all cases, entropy remained below the geometric bound, which defines the maximum-entropy distribution at fixed mean spike count [10] (Fig. S4A).

**Fig. S3.**
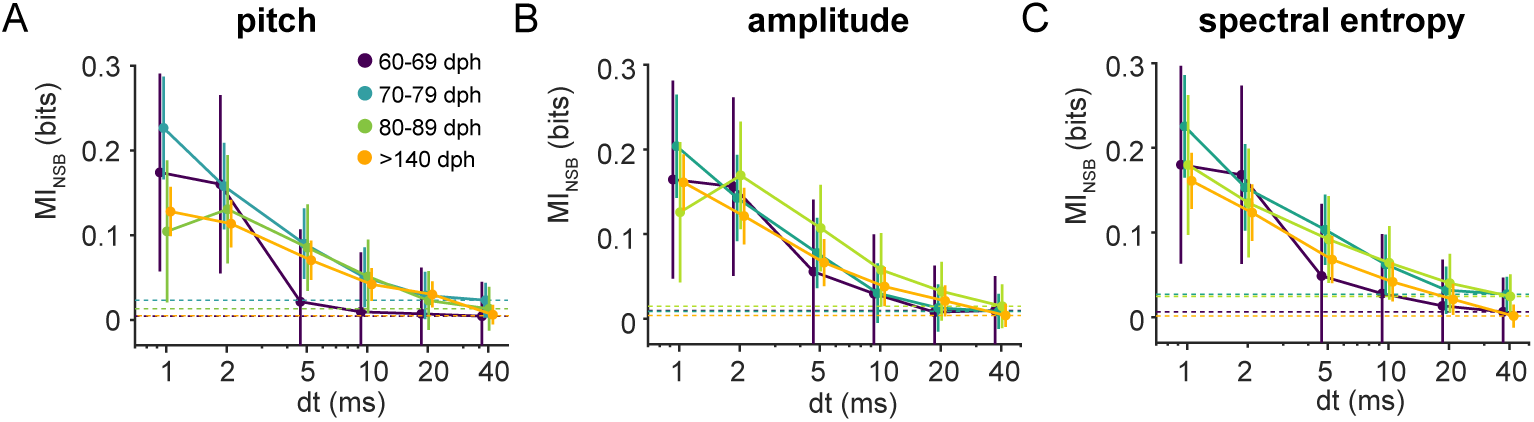
A precise timing code for vocal behavior is observed across all age groups. Mutual information (MI) between premotor spike patterns (40 ms window) and acoustic features was estimated at varying temporal resolutions (*dt*) using the NSB method (see “Methods”). **A–C)** MI values for **(A)** pitch, **(B)** amplitude, and **(C)** spectral entropy. Colors indicate age groups: early juvenile (60–69 dph, purple; 70–79 dph, teal), late juvenile (80–89 dph, green), and adult (*>*140 dph, orange). Across all ages and features, MI significantly increases as temporal resolution becomes finer (*dt →* 1 ms). Conversely, MI at the timescale of the full window (*dt* = 40 ms, equivalent to gross spike count) is negligible and close to the chance level (dashed lines), indicating that firing rate variations do not encode acoustic features. Error bars represent *±*1 SD. Adult data (*>*140 dph) reanalyzed from [9].

In contrast, when we resolve spike patterns at millisecond precision (*dt* = 1 ms), the relevant vocabulary is no longer spike *counts* but spike *timing patterns* (spike words across 1-ms bins). Although raw entropy again grows with mean spike count (Fig. S4C), it now falls substantially below the refractory Poisson reference, indicating that fine-timescale spike trains are more constrained than expected from independent firing given the rate (Fig. S4C). Critically, this departure from Poisson-like statistics becomes stronger with development: the normalized entropy (ratio of raw entropy and rate-matched Poisson entropy) decreases systematically across age at *dt* = 1 ms (Fig. S4D), whereas it remains near unity at *dt* = 40 ms (Fig. S4B). Together, these comparisons show that developmental changes are modest at the spike-count level but pronounced at the millisecond timescale, consistent with increasing temporal structure (and a progressively more selective spike-word vocabulary) during learning.

### Entropy partitioning and variance analysis

To validate the reliability of our entropy estimates, we sought to verify that the variance in our results reflects true fluctuations in neural pattern variability rather than statistical artifacts arising from limited sample sizes. Specifically, we partitioned the unconditional entropy into two distinct components: one determined by the probability of the most frequent state (typically silence or the most common spike word) and one determined by the internal entropy of all remaining states. By analytically decomposing the total variance of the entropy estimate into terms associated with these components, we can determine if our error bars are dominated by simple sampling noise (uncertainty in the frequency of the most common event) or if they genuinely capture the uncertainty in the distribution of rare spike patterns.

**Fig. S4.**
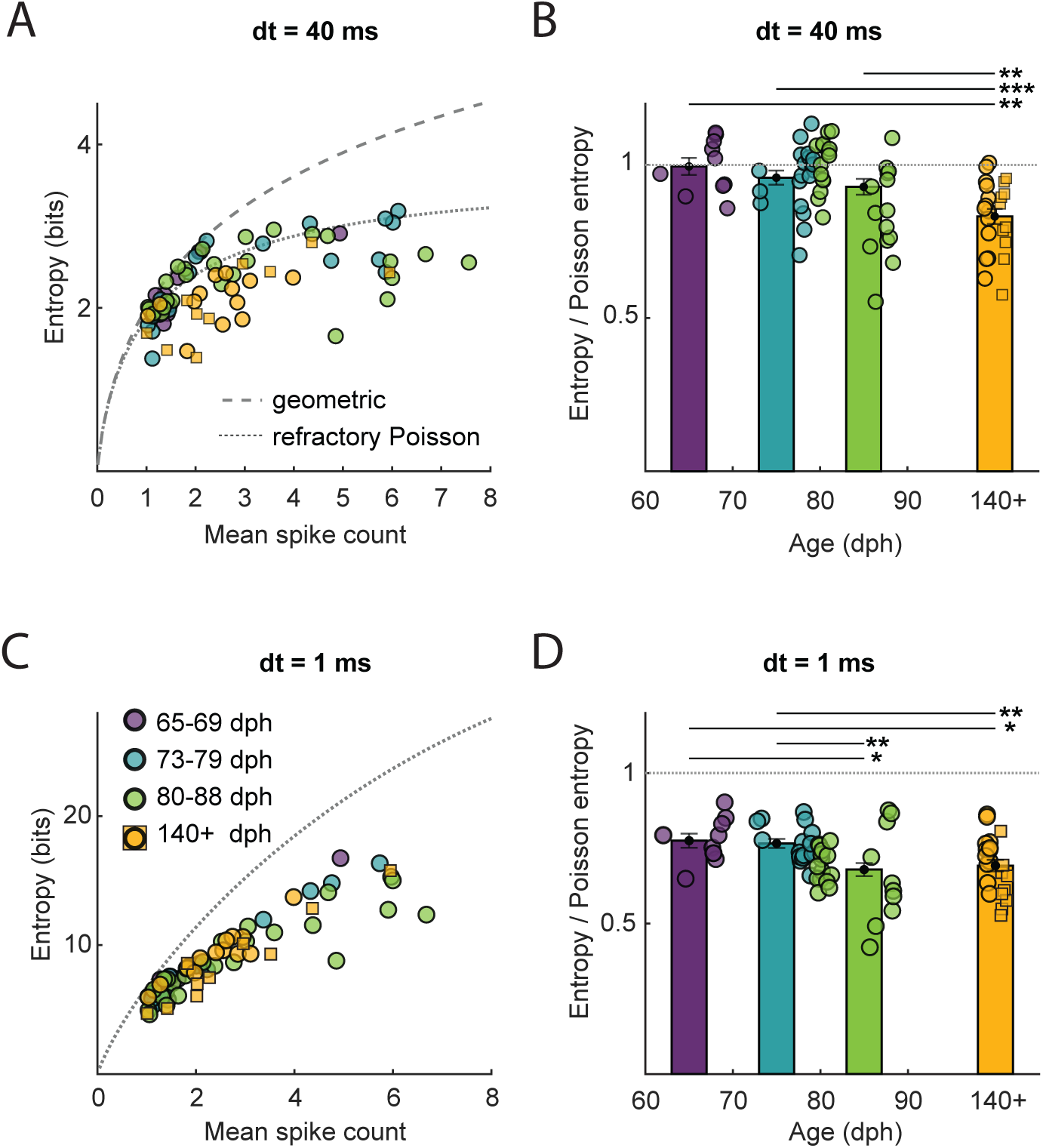
Entropy of RA spike trains at coarse (*dt* = 40 ms) and fine (*dt* = 1 ms) temporal resolutions. Entropy was calculated using the NSB estimator (see “Methods”) for individual “cases” (neuron-syllable pairs) shown as points colored by age: 65–69 dph (purple), 73–79 dph (teal), 80–88 dph (green), and *>*140 dph (orange). **A)** Raw entropy of spike-count distributions (*dt* = 40 ms) plotted against mean spike count for each neuron/day. Dashed and solid curves show the expected entropy of a refractory Poisson process and geometric distribution, respectively. Entropy increases monotonically with mean spike count and approaches—but does not reach—the refractory Poisson limit. **B)** Normalized spike-count entropy (ratio of observed entropy to rate-matched Poisson entropy) as a function of age (*dt* = 40 ms). Values remain close to unity (gray dashed line) across development, indicating that spike-count variability is well explained by an independent-firing (refractory Poisson) model given the mean rate. Small but significant age differences suggest a modest decline in departure from Poisson-like firing. **C)** Raw entropy of millisecond-resolved spike-word distributions (*dt* = 1 ms) versus mean spike count. Entropy grows with increasing mean spike count but remains substantially below the refractory Poisson maximum, reflecting temporal structure beyond rate statistics. **D)** Normalized spike-word entropy at *dt* = 1 ms across ages are much lower than the unity line and declines systematically with age, indicating that fine-timescale spike patterns become less Poisson-like—that is, more temporally structured—as learning progresses. Values for previously reported data in adult birds [5, 9] are shown as square symbols. Bars show group means ± SEM; horizontal lines indicate significant pairwise comparisons (*p *<* 0.05, **p *<* 0.01, **p *<* 0.001).

**Fig. S5.**
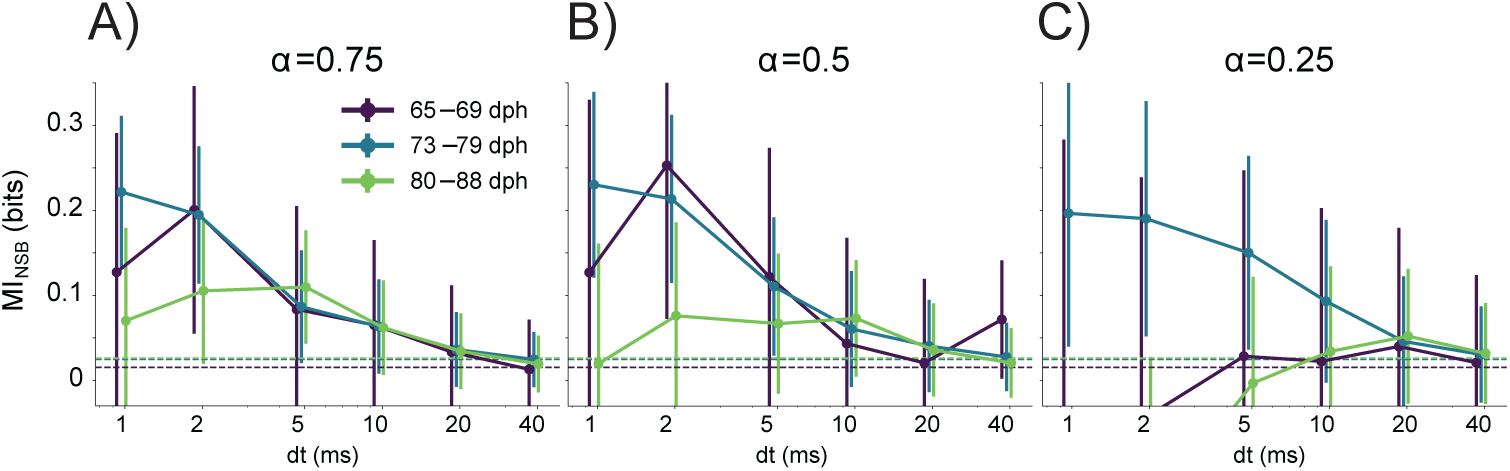
Behavior of the mutual information as a function of the data set size supports the hypothesis of a temporally precise neural code across the sensorimotor period. We plot MI estimates as a function of the discretization scale *dt* of RA activity patterns and the sample set size for different age categories. Each panel correspond to a different fraction *α* of the full dataset of size *N* used for MI estimation. **A**) *α* = 0.75, **B**) *α* = 0.5, **C**) *α* = 0.25. Below *α* = 0.5, MI at high temporal resolution is statistically indistinguishable from zero, suggesting that the evidence of temporally precise neural code in Fig 2 is not an artifact of sample-size dependent estimation biases.

#### Derivation

We start from the expression for the entropy of a disjoint mixture with *X*_1_ and *X*_2_ discrete random variables over disjoint alphabets. We set

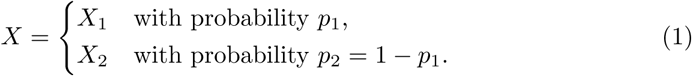

The alphabet for variable *X*_1_ comprises here only a single element, the most common word, with corresponding frequency of occurrence *p*_1_. The alphabet for variable *X*_2_ represents the set of all other words, with the overall frequency of occurrence *p*_2_. We introduce the indicator function *θ*(*X*) such that

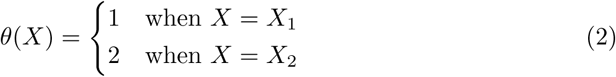

Starting from the chain rule of conditional entropy [11], we write

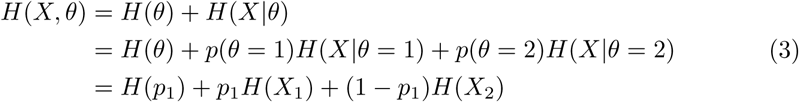

with *H*(*p*_1_) = −*p*_1_ log *p*_1_ −(1 − *p*_1_) log(1 − *p*_1_) the “entropy of choice” between *X*_1_ and *X*_2_. As the alphabet of *X*_1_ contains a single element, namely the most common word in the distribution, we get *H*(*X*_1_) = 0 and the formula for the partitioned entropy becomes

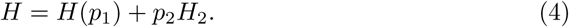

We can then express the variance of the partitioned entropy as:

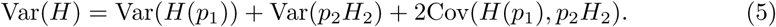

To estimate the variance of the entropy of choice *H*(*p*_1_) we use the “delta method” [12], which states that Var[*f* (*x*)] ≈ Var(*x*) [*f ^′^* (E(*x*))]^2^. This gives

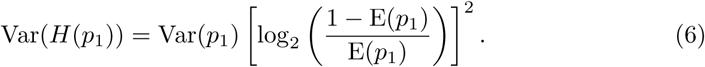

We then use the fact that *p*_2_ and *H*_2_ are independent variables to write

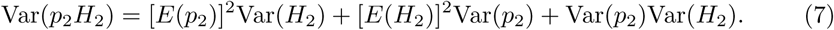

The variance of *H*_2_ is obtained from the variance of the posterior distribution returned by the NSB estimator. The frequency of occurrence *p*_2_ for the set of all words that are not the most common word can be written as *p*_2_ = *n*_2_*/N*, where *n*_2_ is the number of times any word that is not the most common word appeared out of *N* trials. We then use the assumption that *n*_2_ follows the binomial distribution with variance *Np*_2_(1−*p*_2_) to get Var(*p*_2_) = *p*_2_(1 − *p*_2_)*/N* = Var(*p*_1_). Finally, the last term in Eq. [5] can be expressed as

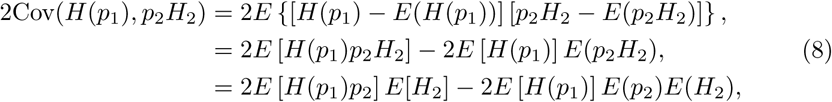

where we used the fact that *p*_2_ and *H*_2_ are independent variables. Estimating the expectations E [*H*(*p*_1_)*p*_2_] and E [*H*(*p*_1_)] up to the second moment in *p*_2_, one obtains:

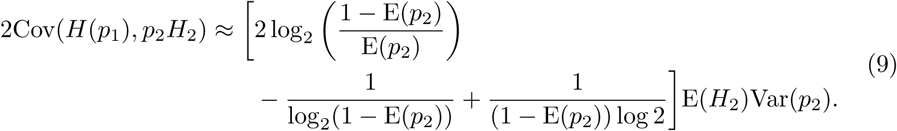

Eq. [5], together with Eqs. [6], [7] and [9] fully specify the error made on the estimate of the partitioned entropy.

#### Entropy partitioning results

The contributions from each term in Eq. [5] are plotted in Fig. S6. We find that at fine temporal resolutions (*dt* → 1 ms), the total variance is dominated by the term Var(*p*_2_*H*_2_) (orange curve), which corresponds to the uncertainty in the entropy of the spike patterns, rather than the variance associated with the partition size *p*_1_ (blue curve). This indicates that the uncertainty in our entropy estimates is primarily driven by the intrinsic variability of the spike trains within the partition, rather than by sampling artifacts related to the relative frequency of the most common word. Consequently, our variance estimates accurately reflect the difficulty of estimating the entropy of the neural code itself, confirming that the NSB estimator is operating effectively even in the regime of sparse data.

**Fig. S6.**
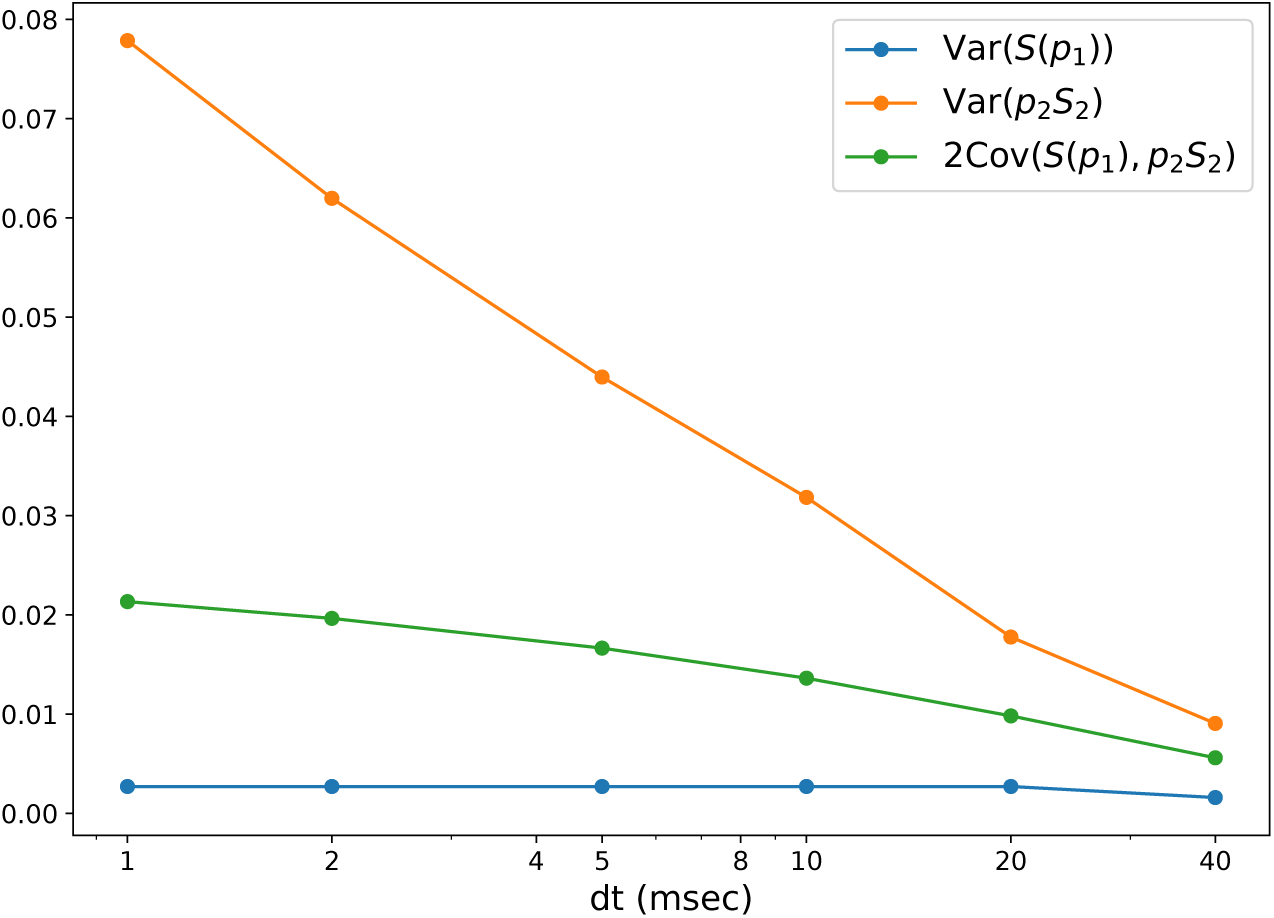
Decomposition of entropy variance across temporal resolutions. The total variance of the partitioned unconditional entropy (Eq. 5) was decomposed to identify the dominant sources of variability in our estimates. Curves show the contributions of the three variance terms as a function of the binning size *dt* for a representative syllable: 1) variance due to fluctuations in partition size (Var(*S*(*p*_1_)), blue), 2) variance due to fluctuations in partition entropy (Var(*p*_2_*S*_2_), orange), and 3) the covariance between them (2Cov(*S*(*p*_1_)*, p*_2_*S*_2_), green). At fine temporal scales (*dt* = 1 ms), the variance is dominated by the entropy term (orange), whereas partition size fluctuations (blue) contribute negligibly. This indicates that uncertainty in our entropy estimates is primarily driven by the variability of spike patterns within partitions, rather than by variations in the number of samples per partition.

### Stationarity tests and data stability

As detailed in the Materials and Methods, we assessed the dataset for significant temporal correlations or non-stationarities that could bias the NSB entropy estimates. To do this, we compared estimates computed from the first and second halves of the recording session (which exposes slow temporal drift, such as changes in firing rate) against estimates computed from interleaved even and odd trials (which serves as a randomized control for sampling variance), as shown in Fig. S7.

**Fig. S7.**
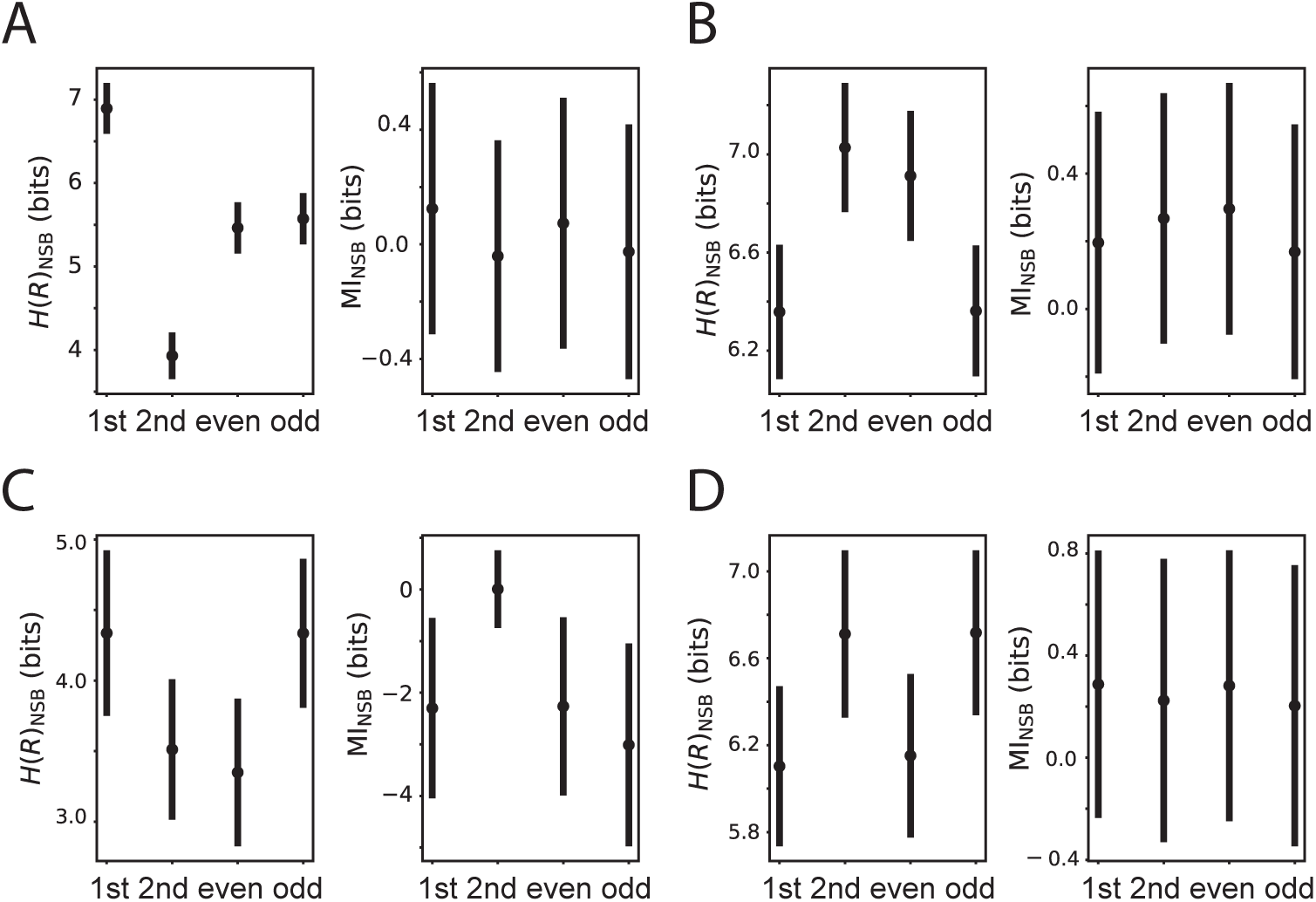
Tests for temporal stability and stationarity in neural recordings. Unconditional entropy (*H*_NSB_(*R*), left subpanels) and MI (MI_NSB_, right subpanels) were estimated at *dt* = 1 ms for single syllables. Panels display data for: **A)** Bird 1 at 68 dph, **B)** Bird 1 at 73 dph, **C)** Bird 2 at 81 dph, and **D)** Bird 2 at 153 dph. Estimates were computed for the first versus second halves of the recording session (“1st” vs. “2nd”) to detect temporal drift and compared against interleaved trials (“even” vs. “odd”) serving as a randomized control. **Note:** These specific examples were chosen for display because they exhibited the greatest potential non-stationarities found in the entire dataset (visible as shifts in unconditional entropy in A and B). Crucially, despite these fluctuations in overall firing statistics, the corresponding MI estimates remain statistically stable across splits. Furthermore, the magnitude of these discrepancies is small relative to the large differences in information observed across the full developmental trajectory.

The examples presented in Fig. S7 were specifically selected because they exhibited the greatest potential non-stationarities (e.g., shifts in unconditional entropy, H(R)) found across the entire dataset. We find that while unconditional entropy may drift slightly in two examples (Fig. S7A-B, left), the resulting MI estimates remain statistically stable across splits. Furthermore, the magnitude of these differences (in bits) is relatively small compared to the overall range of entropy values observed across the developmental dataset. Therefore, we conclude that temporal correlations in our data are minimal and do not introduce systematic bias into our entropy and MI estimates.

**Fig. S8.**
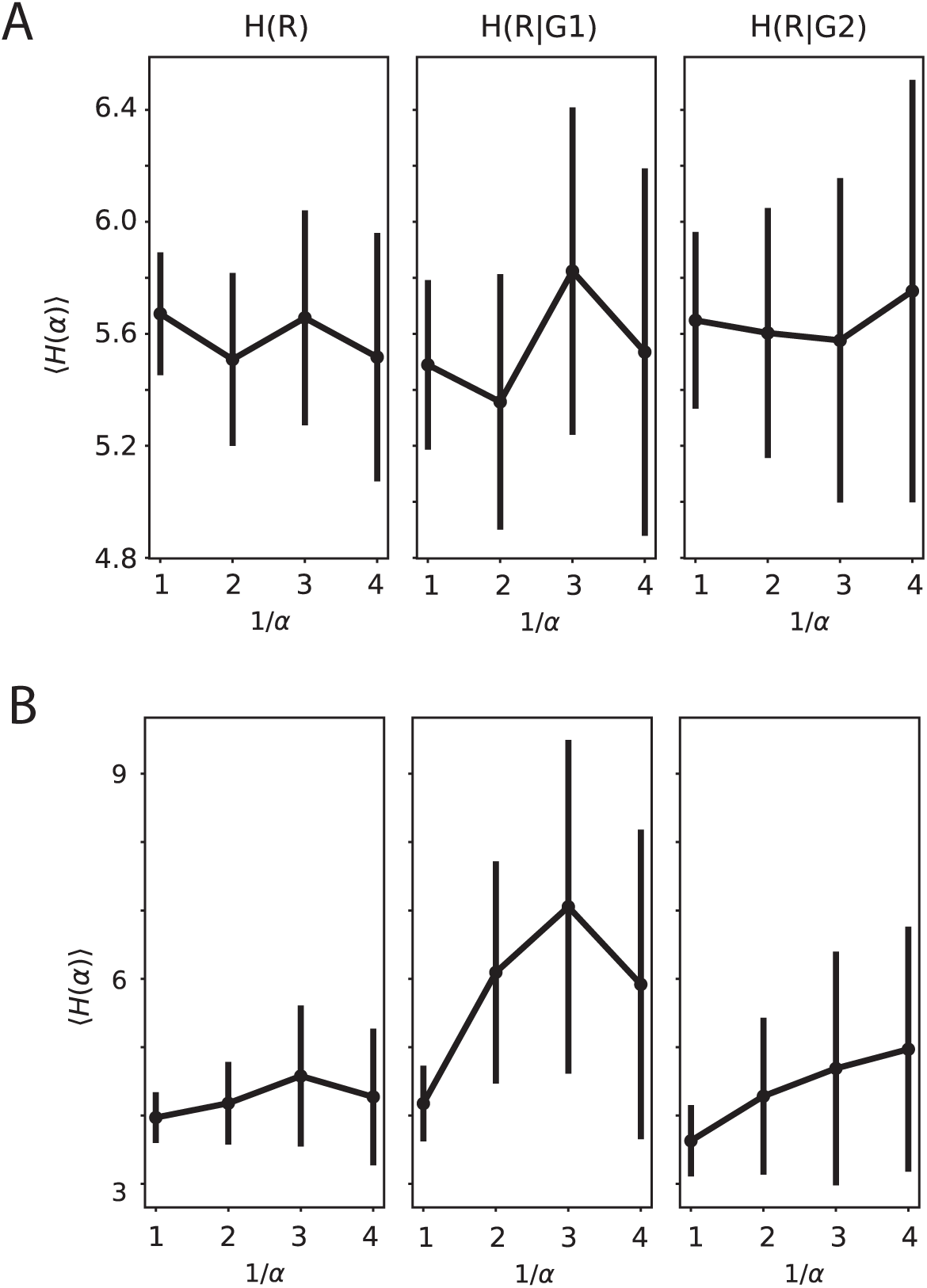
Verification of sample size sufficiency for entropy estimation. Finite data size scaling of the unconditional entropy *H*(*R*) and conditional entropies *H*(*R|G*_1_) and *H*(*R|G*_2_) (conditioned on acoustic groups). Estimates are plotted against the inverse data fraction 1*/α*, where *α* represents the proportion of randomly selected trials used. **A)** Bird 1 (68 dph). **B)** Bird 2 (81 dph). In both examples, entropy estimates remain stable as the data fraction decreases (increasing 1*/α*), and sub-sampled estimates fall within the error bars of the full dataset (1*/α* = 1). This confirms that the full dataset size is sufficient to reach the asymptotic regime where sampling bias is minimal. Syllables analyzed correspond to those in Fig. S7 (Panels A and C). All estimates use a bin size of *dt* = 1 ms; convergence is faster for coarser time bins (*dt >* 1 ms).

### Finite data bias

We estimated the entropy *S*(*α*) for *αN* number of trials, with *α <* 1 and *N* the total number of trials recorded, by randomly selecting trials and averaging over 10 realizations of this subsampling to yield the average entropy estimate ⟨*S*(*α*)⟩ for each data fraction *α*. Results of the data fraction analysis on the unconditional and conditional entropy estimates can be found in Fig. S8.

As all entropy estimates for different data fractions agreed within error bars, we could confirm the absence of an empirical sample-size-dependent bias.

